# Muon: multimodal omics analysis framework

**DOI:** 10.1101/2021.06.01.445670

**Authors:** Danila Bredikhin, Ilia Kats, Oliver Stegle

## Abstract

Advances in multi-omics technologies have led to an explosion of multimodal datasets to address questions ranging from basic biology to translation. While these rich data provide major opportunities for discovery, they also come with data management and analysis challenges, thus motivating the development of tailored computational solutions to deal with multi-omics data.

Here, we present a data standard and an analysis framework for multi-omics — *MUON* — designed to organise, analyse, visualise, and exchange multimodal data. MUON stores multimodal data in an efficient yet flexible data structure, supporting an arbitrary number of omics layers. The MUON data structure is interoperable with existing community standards for single omics, and it provides easy access to both data from individual omics as well as multimodal dataviews. Building on this data infrastructure, MUON enables a versatile range of analyses, from data preprocessing, the construction of multi-omics containers to flexible multi-omics alignment.

## Introduction

Multi-omics designs, that is the simultaneous profiling of multiple omics or other modalities for the same sample or cells, have recently gained traction across different biological domains. Multi-omics approaches have been applied to enable new insights in basic biology and translational research [1,2].

On the one hand, the emerging multi-omics datasets result in novel opportunities for advanced analysis and biological discovery [3]. Critically however, multi-omics experiments and assays pose considerable computational challenges, both concerning the management and processing as well as the integration of such data [4,5]. Major challenges include efficient storage, indexing and seamless access of high-volume datasets from disk, the ability to keep track and link biological and technical metadata, and dealing with the dependencies between omics layers or individual features. Additionally, multi-omics datasets need to be converted into specific file formats to satisfy input requirements for different analysis and visualisation tools.

While specialized frameworks for the analysis of different omics data types have been proposed, including for bulk and single-cell RNA-seq [6–9] or epigenetic variation data [10–13], there is a lack of comprehensive solutions that specifically address multi-omics designs. Additionally, there currently exists no open exchange format for sharing multi-omics datasets that is accessible from different programming languages. The few existing solutions for multi-omics data (Seurat [8], MultiAssayExperiment [14]) are confined to the R programming language ecosystem, and require loading the full dataset into the working memory, which precludes dealing with larger datasets.

To address this, we here present MUON (**mu**ltimodal **o**mics a**n**alysis), an analysis framework that is designed from the ground-up to organise, analyse, visualise, and exchange multimodal data. MUON is implemented in Python and comes with an extensive toolbox to flexibly construct, manipulate and analyse multi-omics datasets. At the core of the framework is MuData, an open data structure standard, which is compatible with and extends previous data formats for single omics [9,15]. MuData files can be seamlessly accessed from different programming languages, including Python [16], R [17], and Julia [18]. We illustrate MUON in the context of different vignettes of its application, including analysis of combined gene expression and chromatin accessibility assays as well as gene expression and epitope profiling.

## Results

### MuData: a cross-platform multimodal omics data container

At the core of MUON is MuData (***mu****ltimodal* ***data***) — an open data structure for multimodal datasets. MuData handles multimodal datasets as containers of unimodal data. This hierarchical data model generalizes existing matrix-based data formats for single omics, whereby data from each individual omics layer are stored as an AnnData [15] object. (**Figure 1a**,**c**). MuData also provides a coherent structure for storing associated metadata and other side information, both on the level of samples (e.g. cells or individuals) and features (e.g. genes or genomics locations). Metadata tables can either be specific to a single stored data modality, or they can represent joint sample annotations that apply to all modalities stored in a MuData container. In a similar manner, MuData containers can be used to store derived data and analysis outputs, such as cluster labels or an inferred sample embedding (**Figure 1b**).

**Fig. 1:**
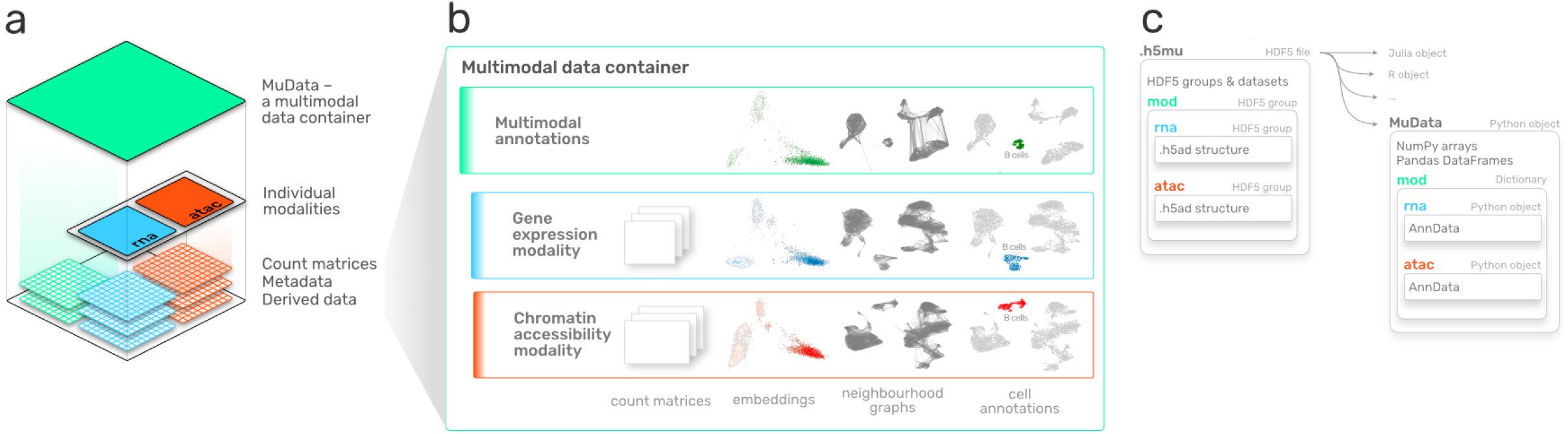
Architecture and content of a multimodal data container (MuData). **a**. Schematic representation of the hierarchical structure of a MuData container. Raw data matrices from multiple modalities together with associated metadata are encapsulated in an array structure. For illustration, blue and red denote RNA-seq and ATAC-seq data modalities; green denotes multimodal annotation or derived data. **b**. Example content of the structure in **a**. Shown are example content of a MuData container, consisting of count matrices, embeddings, neighbourhood graphs and cell annotations for individual modalities (blue, red), as well as derived data from multi-omics analyses (green). **c**. Schematic representation of MUON storage model and its serialization scheme using the HDF5 file format on disk. **Left:** Hierarchy of the storage model, with plates denoting different levels of hierarchy. Arrows signify access schemes of the HDF5 file using various programming languages. **Right:** Representation of the MuData object in Python, with metadata and derived annotations represented as NumPy arrays or Pandas DataFrames, and with individual modalities as AnnData objects.

MuData objects are serialised to HDF5 [19] files by default — the industry standard for storing hierarchical data. Individual omics layers are serialised using the existing AnnData serialisation format, thus permitting direct access to single omics using existing toolchains that build this data standard (**Figure 1c**). Basic access to MuData objects is possible from all major programming environments that support access to HDF5 array objects. Additionally, MUON comes with dedicated libraries to create, read and write MuData objects from Python, R, and Julia. These tools facilitate the exchange of multi-omics data across platforms, and ensure consistent file format definitions.

### MUON: a framework for multimodal omics data

The MUON framework allows for managing, processing, and visualising multi-omics data using the MuData containers. Existing workflows developed for single-omics can be reused and applied to the contents of a multi-omics container. For example, individual modalities of the simultaneous gene expression and chromatin accessibility profiling [20] can be processed using existing RNA [21] and ATAC [22] workflows. In this manner, canonical processing steps, including quality control, sample filtering, data normalization and the selection of features for analysis can be transferred from single-cell omics analysis (**Figure 2a**).

**Fig. 2:**
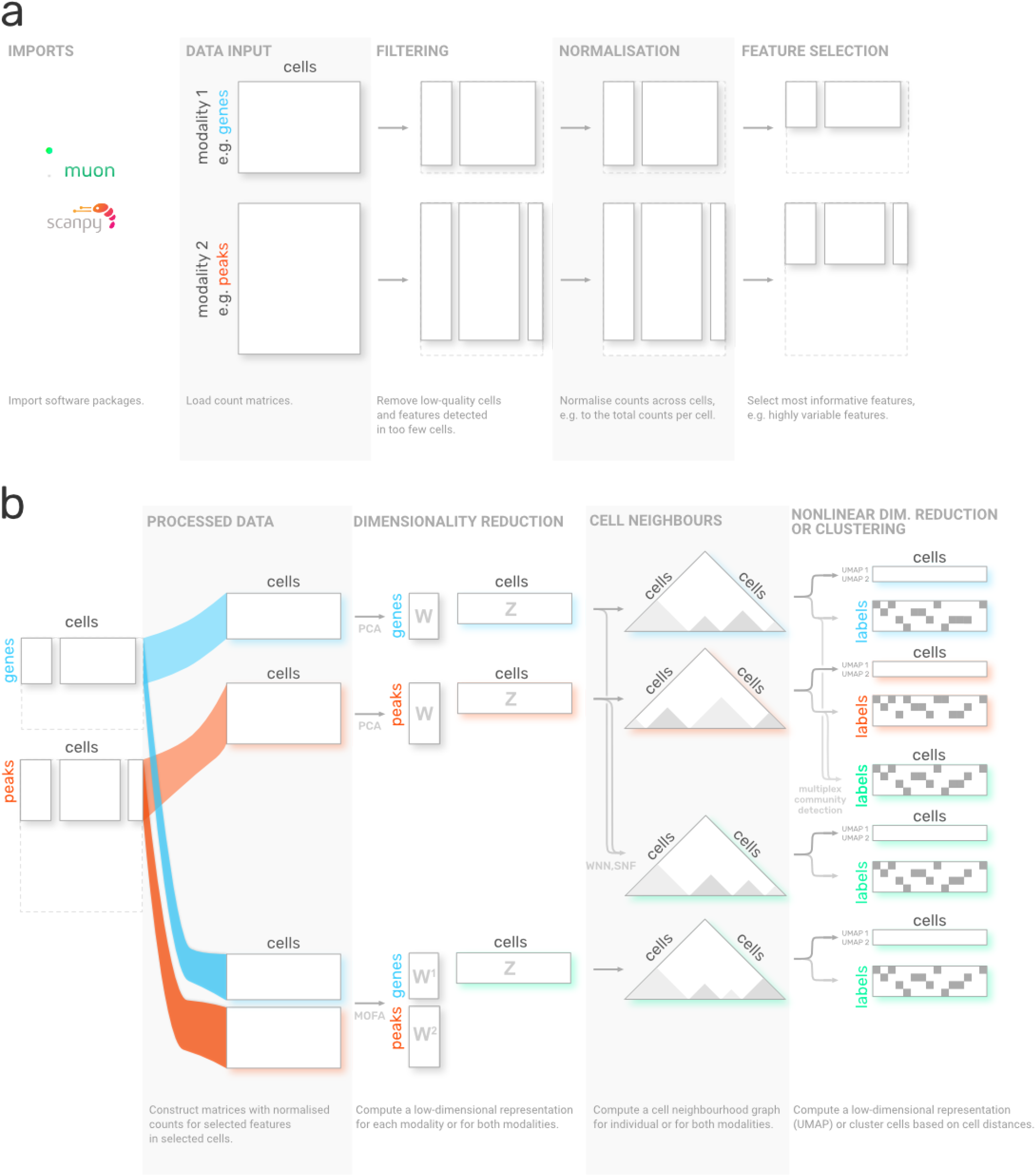
Example multi-omics analysis workflows implemented using MUON. **a**. Construction and processing of individual modalities of a multi-omics scRNA & scATAC dataset. Processing steps for individual omics from left to right. Rectangles denote count matrices following each processing step, which are stored in a shared MUON data container. MUON provides processing functionalities for a wide range of single-omics, including RNA-seq, ATAC-seq, CITE-seq. Existing workflows and methods can be utilized, including those implemented in scanpy. Respective analysis steps are described below each step. **b**. Alternative workflows for integrating multiple omics for latent space inference and clustering. MUON enables combining alternative analysis steps to define tailored multi-omics data integrations. Shown are canonical workflows from left to right: dimensionality reduction, definition of cell neighbourhood graphs, followed by either nonlinear estimation of cell embeddings or clustering. Triangles represent cell-cell distance matrices, with shading corresponding to cell similarity. Green colour signifies steps that combine information from multiple modalities; steps based on individual modalities only are marked with blue (RNA) or red (ATAC) respectively. The outputs of the respective workflows (right) are from top to bottom: UMAP space (i) and cell labels (ii) based on RNA or alternatively based on ATAC modality (iii, iv), cell labels based on two cell neighbour graphs from individual modalities (v), UMAP space and cell labels based on WNN output (vi, vii), UMAP space and cell labels based on MOFA output (viii, ix).

The integration of multiple modalities within a MuData container facilitates the definition of multi-omics analysis workflows, allowing to flexibly combine alternative processing steps (from left to right in **Figure 2b**). For example, single-omics dimensionality reduction methods such as principal component analysis or factor analysis [23–26] can be used to separately process scRNA-seq & scATAC-seq count matrices. Additionally, MUON comes with interfaces to multi-omics analysis methods that jointly process multiple modalities, including multi-omics factor analysis [27,28] (MOFA) to obtain lower-dimensional representations, and weighted nearest neighbours [29] (WNN) to calculate multimodal neighbours. Once the results from either dimensionality reduction strategy are stored in a MUON container, they can be used as input for defining cell neighbour graphs. This graph can be either estimated from individual omics modalities, from a multi-omics representation (e.g. as obtained from MOFA), or by fusing two single-omics neighbour representations (e.g. using methods such as similarity network fusion, SNF [30], or WNN [29]).

Finally, the latent or neighbourhood representations can serve as a starting point for downstream analysis and interpretation. For example, UMAP [31] can be directly applied to cell neighbourhood graphs to generate nonlinear embeddings of cells. Similarly, the alternative cell neighbourhood graphs can be used as input for identifying connected components and thereby putative cell types (e.g. using multiplex community detection techniques [32]).

The flexibility to choose and control individual processing steps in MUON makes it possible to compose tailored workflows for a particular dataset.

### Application of MUON to single-cell multi-omics data

To illustrate MUON, we considered data from simultaneous scRNA-seq & scATAC-seq profiling of peripheral blood mononuclear cells (PBMCs), which were generated using the Chromium Single Cell Multiome ATAC + Gene Expression protocol by 10x Genomics [20]. Features in the RNA modality correspond to the expression level of genes, whereas the ATAC modality encodes accessible genomic loci as peaks. MUON supports the application of alternative dimensionality reduction strategies (**Figure 2**). For example, multi-omics factors analysis [28] yields a lower dimensional representation, including factors that capture variation of individual omics or shared variability (**Figure S1**), which in turn can be interpreted on the level of individual features (**Figure 3a**). Here, the factors that explain the largest fraction of variance in PBMCs capture canonical biological differences, such as the myeloid – lymphoid axis and cytotoxicity (**Figure 3a, left**). These factors capture both variation in mRNA abundance and chromatin accessibility, e.g. as CD3E expression and BCL11B promoter accessibility, which are characteristic for T cells [33,34] (**Figure 3a, right**).

**Fig. 3:**
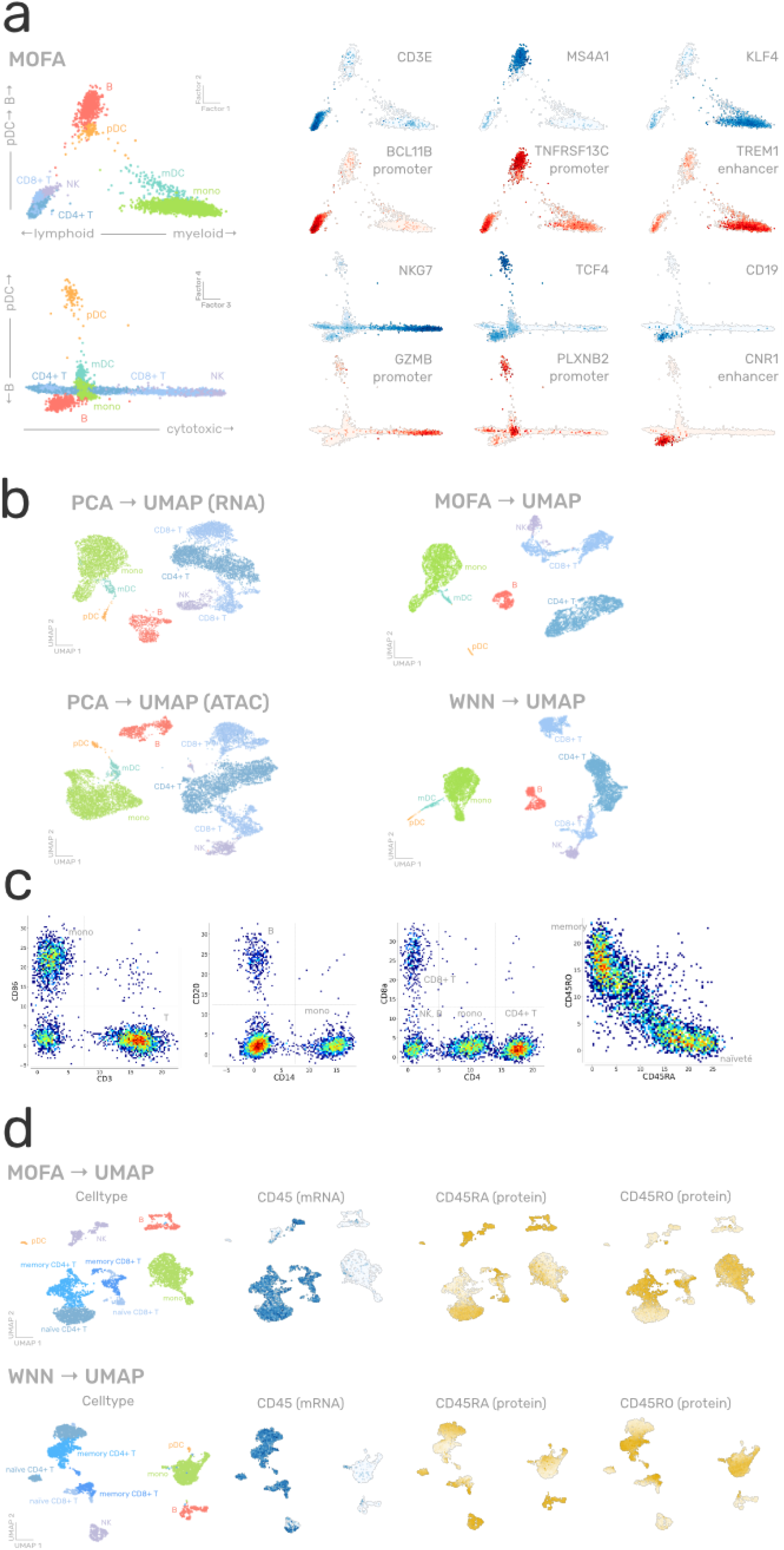
Single-cell multi-omics datasets processed and visualised using MUON. **a**. MOFA factors estimated from simultaneous scRNA-seq and scATAC-seq profiling of PBMCs, with cells coloured by either left: coarse-grained cell type; or right: gene expression (in blue) and peak accessibility (in red). Displayed genes and peaks are selected to represent cell type-specific variability along factor axes. **b**. UMAP latent space for the same dataset as in **a**, constructed from left: principal components for individual modalities; or right: MOFA factors and WNN cell neighbourhood graph. Cells are coloured by coarse-grained cell type. **c**. Examples of individual feature values of protein abundance in the CITE-seq profiling of PBMCs after applying dsb normalization. Colours correspond to the relative local density of cells with red for high density and blue for low density. **d**. UMAP latent space for the same dataset as in **c**, constructed from MOFA factors (top) and WNN cell neighbourhood graph (bottom). Cells are coloured by their coarse-grained cell type or feature values (blue for gene expression, yellow for protein abundance).

A two-dimensional latent space recapitulating the structure of the data is commonly used for visualising cell type composition, cell-level covariates, or feature counts. For this, it is important that MUON allows to generate, store, and operate with multiple different embeddings constructed for individual modalities (**Figure 3b, left**) or jointly for both modalities based, for instance, on the MOFA factors or the WNN graph (**Figure 3b, right**). Such visualisations can be generated from MuData objects without loading all the data into memory.

As a second example, we considered CITE-seq [35] data, which comprises gene expression and epitope abundance information in the same cells. To process the latter, specialized normalisation strategies for denoising and scaling [36] are available. Normalised protein counts can then be used to define cell types, akin to gating in flow cytometry [37] (**Figure 3c**). Once the count matrices are processed, these can be integrated using alternative multimodal options (**Figure 2**). For instance, using both modalities for cell type annotation as well as for dimensionality reduction allows to attribute the distinction between naïve and memory T cells to the abundance of CD45 isoforms RA and RO at the protein level (**Figure 3d**).

## Discussion

Multimodal omics designs are increasingly accessible, allowing for characterising and integrating different dimensions of cellular variation, including gene expression, DNA methylation, chromatin accessibility, and protein abundance [3,38,39]. MUON directly addresses the computational needs posed by such multi-omics designs, including data processing, analysis, interpretation, and sharing (**Figure 1**). Designed for the Python ecosystem, MUON operates on MuData objects that build on community standards for single-omics analysis [9]. Serialization to HDF5 makes MuData objects accessible to other programming languages, including R and Julia.

MUON is designed in a modular fashion, which means that existing methods and tools for processing individual omics can be reused to design more complex analysis workflows (**Figure 2**,**3**). At the same time, the software facilitates combining single-omics analysis methods with a growing spectrum of multi-omics integration strategies to define novel multi-omics workflows.

Looking ahead, MUON will be a robust platform to build upon and support future developments. On the one hand, handling novel assays for multi-omics that are emerging can be integrated. For example, mRNA and proteins can be assayed together not only with CITE-seq [35] but also with QuRIE-seq [40] or INs-seq [41]. Moreover, trimodal assays such as scNMT-seq [42] or TEA-seq [43] allow to generate data beyond just two modalities and can be handled with MUON, which is designed to manage an arbitrary number of modalities. On the other hand, the complexity of experimental designs is rapidly increasing [44,45]. Already, MUON can take additional covariates into account during multimodal integration, for example to perform temporally aware factor analysis [46]. Future development of MUON will include incorporating additional relationships in MuData, for example to explicitly model the dependencies between feature sets across omics, or to account for dependencies between multiple sets of multi-omics experiments.

## Methods

### Software availability

The code is made available at https://github.com/gtca/muon under the BSD3 license, and its documentation can be accessed at https://muon.readthedocs.io/. The source code for Julia and R libraries can be accessed at https://github.com/gtca/Muon.jl and https://github.com/gtca/muon.r.

### Implementation of muon

Muon has been implemented in the Python programming language and builds on a number of existing numerical and scientific open-source libraries, in particular NumPy [47], Scipy [48], Sklearn [49], Pandas [50], h5py [51], AnnData [15] and Scanpy [9] for omics data handling, MOFA+ [28] for multimodal data integration and matplotlib [52] and seaborn [53] for data visualisation. The weighted nearest neighbours (WNN) method has been implemented following [29] describing the original method and [43] describing its generalisation to an arbitrary number of modalities.

### Comparison of MUON with alternative analysis frameworks

MUON takes inspiration and builds on concepts from Scanpy [9]. In fact, the software incorporates ideas and extends it in a modular fashion, similar to the existing practice in the Bioconductor community [54].

|  | **muon** | **scanpy** [9] | **Seurat** [8] | **MultiAssay Experiment** [14] |
| --- | --- | --- | --- | --- |
| Main programming environment | Python | Python | R | R |
| Working with data without loading it all into memory | Yes | Yes | No | No |
| Serialization accessible from multiple languages | Yes (.h5mu) | Yes (.h5ad) | No (.rds) | No (.rds) |
| Native support for I/O operations | Python, Julia, R | Python | R | R |
| Support for multiple modalities | Yes | No | Yes | Yes |
| Support for data missing in some modalities | Yes | NA | No | Yes |
| Support for multimodal embeddings | Yes | NA | No | No |

### Processing gene expression and chromatin accessibility data

Single-cell multiome ATAC + gene expression demonstration data for peripheral blood mononuclear cells (PBMCs) from a healthy donor with granulocytes removed through cell sorting processed with ARC 1.0.0 pipeline was provided by 10X Genomics (https://support.10xgenomics.com/single-cell-multiome-atac-gex/datasets).

Log-normalisation was used for both gene and peak counts, and respective values for highly variable features scaled and centered to zero mean and unit variance were then used as input to discussed algorithms such as PCA, as implemented in scikit-learn [49] and scanpy [9], or MOFA+ [28]. Differentially expressed genes and differentially accessible peaks were identified with respective functionality in scanpy and were used to compile gene lists for cell type identification.

The respective vignettes are available at https://muon-tutorials.readthedocs.io/en/latest/single-cell-rna-atac.

### Processing CITE-seq data

CITE-seq data for PBMCs from a healthy donor was provided by 10X Genomics (https://support.10xgenomics.com/single-cell-gene-expression/datasets/3.0.2/5k_pbmc_protein_v3). Log-normalisation was used for gene counts, and dsb [36] was used to denoise and scale protein counts. Respective values for highly variable features scaled and centered to zero mean and unit variance were then used as input to discussed algorithms. The respective vignettes are available at https://muon-tutorials.readthedocs.io/en/latest/cite-seq.

## Acknowledgements

We are grateful to the members of the Stegle and Theis labs for the discussions on the MuData and MUON design. D.B. is supported by the EMBL International PhD Programme and a Darwin Trust fellowship.

---

**Fig. S1:**
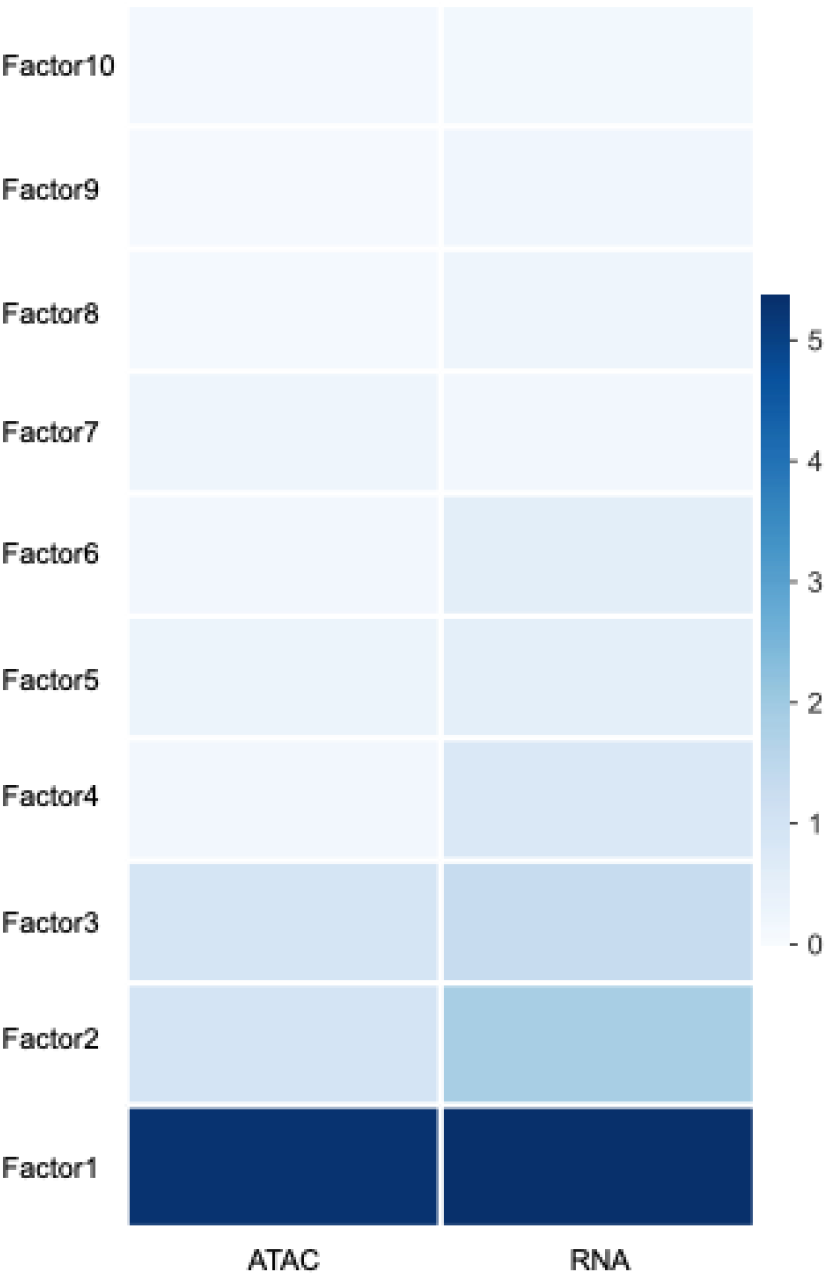
Variance explained by each of 10 MOFA factors in each modality of simultaneous scRNA-seq and scATAC-seq profiling of PBMCs. Colour denotes the percentage of variance explained.

